# Vector-enabled metagenomics reveals the first detection of the geminivirus beet curly top Iran virus in Europe

**DOI:** 10.64898/2026.02.18.706615

**Authors:** Laura Miozzi, Silvia Rotunno, Fulco Frascati, Monica Marra, Francesco Nugnes, Umberto Bernardo, Daniele Marian, Sofia Bertacca, Massimiliano Ballardini, Gian Paolo Accotto, Anna Maria Vaira, Emanuela Noris

## Abstract

2.

Geminiviruses are among the most threatening emerging insect-borne viruses and are responsible for serious outbreaks. Climate change could further exacerbate their impact on crops, highlighting the need for new diagnostic approaches to manage potentially dangerous situations. Vector-Enabled Metagenomics (VEM) exploits the natural ability of highly mobile insects to accumulate viruses acquired from plants over time and space within an ecosystem; this approach is effectual in monitoring the presence of new invasive and indigenous viruses in large areas. Geminiviruses have circular single-stranded DNA genomes that can be readily targeted by Rolling Circle Amplification (RCA). The combination of RCA and VEM largely increases the chances of detecting geminiviruses. This approach enabled us to identify the becurtovirus beet curly top Iran virus (BCTIV, *Becurtovirus betae*) in insects collected in Europe. BCTIV is a major pathogen of sugar beet but can also infect plants of other families; it is transmitted by cicadellids and has been so far detected only in Iran and Anatolia (Turkey). We also show that two cucurbit species, watermelon (*Citrullus lanatus*) and zucchini (*Cucurbita pepo*) are both natural and experimental hosts for BCTIV.

**Impact statement:** Virus infections account for almost 50% of emerging plant diseases globally and may produce high crop losses, resulting in huge economic and social impact worldwide. Geminiviruses, threatening both monocot and dicot plants, represent high risk for both staple food and industrial crops. A peculiar diagnostic approach combining a specific geminivirus enrichment reaction, the monitoring of the virome of highly mobile insects within agricultural areas and the sensitivity of high throughput sequencing (HTS) was effective in producing a first alert for a new polyphagous virus. The reduced cost of HTS methods further raises interest in this approach, making it suitable as a first step for monitoring large areas.

**Data summary:** The authors confirm all supporting data, code and protocols have been provided within the article or through supplementary data files.

## 5. INTRODUCTION

The world’s staple and nutritionally important food crops are increasingly affected by virus disease pandemics and epidemics which significantly reduce their yields and quality. This situation is progressively more alarming due to the growing food requirements for the human population and global warming issues [1]. Emerging infectious diseases are generally caused by pathogens which invade new geographical areas mainly due to the spread of their natural vectors, or attack new species with increased incidence, gaining winning ecological fitness [2]. The *Geminiviridae* family, listing fourteen Genera [3], includes viruses with single-stranded circular DNA (ssDNA) genomes, encapsidated into twinned icosahedral particles. Geminiviruses affect both monocot and dicot plants, including important cereals, vegetables, ornamentals and fiber crops. They are among the most threatening emerging plant viruses and have caused serious outbreaks during the last decades across different parts of the world [4]. The geminivirus beet curly top Iran virus (BCTIV, *Becurtovirus betae*) has a monopartite genome, infects mainly sugar beet (*Beta vulgaris*), but also tomato, pepper, and common bean [5, 6], and is transmitted by *Neoaliturus (Circulifer) haematoceps* (Mulsant & Rey), family Cicadellidae [7]. Up to date, BCTIV has been reported only in the Asian continent, *i*.*e*. in Iran and Anatolia (Turkey).

Metagenomics-based approaches are suitable for a wide epidemiological surveillance of viral pathogens. Specifically, Vector-Enabled Metagenomics (VEM) exploits the natural ability of highly mobile insects to accumulate viruses acquired from plants over time and space within an ecosystem [8]. VEM has already been applied to insect vectors or their predators to identify known and novel plant viruses [9-13].

In this work, we carried out a VEM survey in an intensive vegetable-producing area in Southern Italy, with the aim to analyze the circulating geminivirome. To increase the detection of the geminivirus genomes, a Rolling Circle Amplification (RCA) step was included before VEM analysis [9, 11]. The RCA-VEM procedure was applied to leafhoppers (cicadellids), known to include members that can transmit geminiviruses, but can also be just carriers of viruses after feeding. This strategy enabled us to detect BCTIV for the first time in Italy in insects and in cucurbit plants, providing evidence of the virus presence in the European continent. The ability of BCTIV to infect watermelon and zucchini was also confirmed by inoculation experiments with an infectious agroclone.

## 6. METHODS

### Sample collection and processing

Cicadellids were collected during spring and summer of 2020-2022 in cucurbit and tomato fields and nearby non-cultivated areas located in the Campania region (Southern Italy). Areas of 100 m x 2 m along the fields were surveyed using sweeping nets or entomological aspirators. Cicadellids were transferred into collection tubes and stored in ethanol at -20°C. A total of 74 cicadellid samples (each made of 1 to 8 individuals) were collected from 30 different fields, as described in Table 1.

**Table 1.**
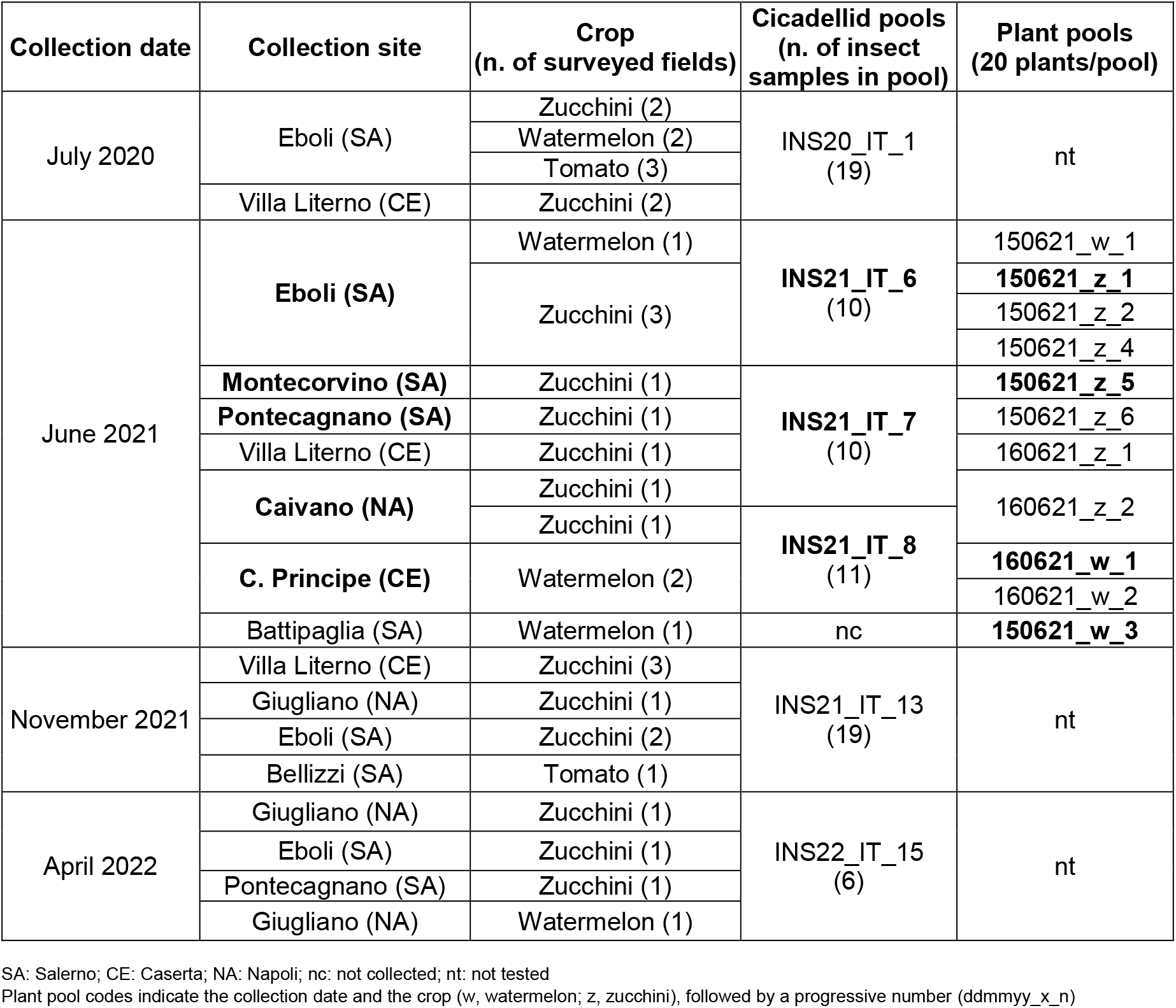
Insect and plant samples analyzed in this study for the presence of beet curly top Iran virus (BCTIV). Collection sites, cicadellid and plant samples where BCTIV-related sequences were identified are indicated in bold.

In the same fields, plant samples consisting of zucchini (*Cucurbita pepo*), watermelon (*Citrullus lanatus*), and tomato (*Solanum lycopersicum*), showing virus-like symptoms were also collected; for each field, 20 representative plants were collected forming a unique plant pool. The plant samples were dried by calcium chloride treatment immediately after collection and stored at -20°C.

### Total nucleic acids (TNA) extraction and RCA amplification

The 74 cicadellid samples were extracted using a CTAB-based method according to [14]. TNA concentration was adjusted to 500 ng/µL, following Nanodrop spectrophotometer (Thermo Scientific) measurement. TNAs were subsequently grouped into 6 pools according to collection date and location (Table 1).

Each insect TNA pool (1.5 µg) was subjected to RCA, using the Sigma TempliPhi Amplification kit (Sigma Aldrich), following manufacturers’ instructions. The reaction was conducted at 30 °C for 30 hours [15]. RCA products were purified using the DNA Clean & Concentrator Kit (Zymo Research) and 1 µg of each purified RCA product was delivered to Novogene service for DNA sequencing with Illumina Novaseq 6000 (2x150bp).

### Cicadellid examination

Cicadellids were morphologically examined based on the available descriptions and diagnostic keys for the genus *Neoaliturus (Circulifer)* [16, 17]. Molecular identification was performed by barcoding on selected BCTIV-positive individual insect DNA, amplifying a fragment of about 680bp of the COI gene (Cytochrome C Oxidase subunit I) using the universal primers LCO/HCO [18], followed by Sanger sequencing. Obtained sequences were compared with those present in the GenBank NCBI (https://www.ncbi.nlm.nih.gov/accessed on 26-Jan-2026) database with BLASTN [19].

### Illumina High throughput Sequencing (HTS) and data analysis

Raw data were checked for quality and adapter contamination with FASTp (v. 0.21.0) [20]. Ribosomal sequences were removed with BBMap tool (v. 38.7, https://sourceforge.net/projects/bbmap/). Reads were assembled into scaffolds using metaSPAdes (v. 3.15.1) [21] with k-mer sizes of 97, 107, 117, and 127; statistics on the assembly were obtained with QUAST (v. 5.2.0) [22]. Redundancy reduction was performed with CAP3 (Version Date 02/10/15) [23]. Only contig sequences longer than 500 bp were retained and used as input for diamond BLASTx (v. 2.0.15; diamond database updated at 13/05/2023) [24]. Results of BLASTx search were visualized with MEGAN (v. 6.25.9) [25]. In order to retrieve full-length geminiviral genomes, clean reads were mapped against the BCTIV reference genome (RefSeq Acc. No. NC_010417.1) using Bowtie2 (v. 2.2.9) [26]. For cicadellid identification, contigs were mapped with Bowtie2 on the COI gene sequences of the Cicadellidae family available in the BOLD database [27] with default parameters.

For phylogenetic analysis, complete genomes and aminoacid sequences of the coat protein (CP) of selected *Becurtovirus* isolates and reference sequences of the genus *Curtovirus* were downloaded from NCBI Virus (https://www.ncbi.nlm.nih.gov/labs/virus/vssi/#/, accessed 20/01/2026). These sequences were multialigned using MUSCLE embedded in MEGA11 software (v. 11.0.11) [28]. The IQTREE online tool (v. 3.0.1, http://iqtree.cibiv.univie.ac.at/) [29] was used for model selection and tree inference. The obtained phylogenetic trees were modified using FigTree (v.1.4.3, https://tree.bio.ed.ac.uk/software/figtree/).

### Validation by end-point PCR and Sanger sequencing

The presence of BCTIV was validated by end-point PCR using the virus-specific primers BCTIV-272-F (5’-CGAAGCTATCCAGCCTTGCT-3’) and BCTIV-831-R (5’-CGATCCACAATAACCCAATG-3’), amplifying a 559 bp fragment of the BCTIV-SIV isolate (GenBank Acc. No. JX082259), spanning the V2 and V1 coding sequences. PCR was performed using Platinum™ II Taq Hot-Start DNA Polymerase kit (Invitrogen), following manufacturer’s instruction, with an annealing temperature of 55 °C. TNA extracted from the field sample 150621_w_3 or from experimentally infected *N. benthamiana* plants was used as positive control. Selected representative amplicons were Sanger-sequenced to verify amplification specificity.

### BCTIV agroinoculation

For infectivity assays, agrobacteria carrying the infectious clone of the BCTIV-SIV isolate [7] were grown for 48 h at 28°C in 50 ml liquid YEB medium containing 100 µg/l kanamycin and 50 µg/l rifampicin, with shaking. Bacteria were pelleted and resuspended in 1.5 ml sterile water. About 50 µL of the suspension were inoculated in the stems of cucurbit seedlings at the first-true leaf stage. Several cultivars of watermelon (4 seedlings of cv. Sugar Baby, 2 of cv. Crimson, 8 of cv. Bontà, and 4 of cv. Sentinel), 14 zucchini seedlings (cv. Genovese), and 4 melon seedlings (*Cucumis melo*, cv. Vendram) were tested in two separate experiments. Seven *N. benthamiana* plants at the 4-leaf stage were inoculated as positive controls. All plants, including non-inoculated controls, were maintained in greenhouse at 25 ± 5 °C, under 16 h light/8 h dark cycle. Plants were evaluated for symptoms and tested by PCR for BCTIV infection 40 days post inoculation (dpi).

## 7. RESULTS AND DISCUSSION

### NGS analysis of insect pools and PCR detection

The high-throughput sequencing of DNAs extracted from the six cicadellid samples generated a total of 837,226,900 reads. Following pre-processing and quality filtering, approximately 99,328,568 to 150,180,006 reads per sample were retained for downstream analysis (Supplementary Table 1). BCTIV-related sequences were identified in the three pools INS21_IT_6, INS21_IT_7 and INS21_IT_8 (Table 1). Specifically, a full-length genome sequence of BCTIV was recovered in pools INS21_IT_7 and INS21_IT_8, showing 99% identity to the BCTIV-SIV isolate genome (Acc. No. JX082259). The INS21_IT_7-derived sequence was submitted to the GenBank database with the accession number PV972859. In the INS21_IT_6 pool, a partial BCTIV genome sequence of 533 nt was recovered, spanning nucleotides 2435-120 of the BCTIV-SIV isolate, including the Large Intergenic Region, and showing 99% identity with the BCTIV-SIV isolate. The detection of a partial BCTIV sequence in this pool could result from a very low concentration of viral target DNA (Supplementary Table 1) or by a technical bias in library construction [30, 31].

Both genome- and CP-based phylogenetic analyses revealed a well-defined BCTIV cluster, clearly distinct from other becurtoviruses (Fig. 1, Supplementary Fig. 1). Within this cluster, two main subgroups are evident: one comprising isolates from central and Southern Iran and another including isolates from Northern and Western Iran and from Turkey. The latter forms a cohesive subclade with isolates from Iran’s Azerbaijan province, near the Turkish border. Notably, the Italian isolate clusters within the central/Southern Iran subgroup, indicating a close genetic relationship with isolates from this region.

**Figure 1.**
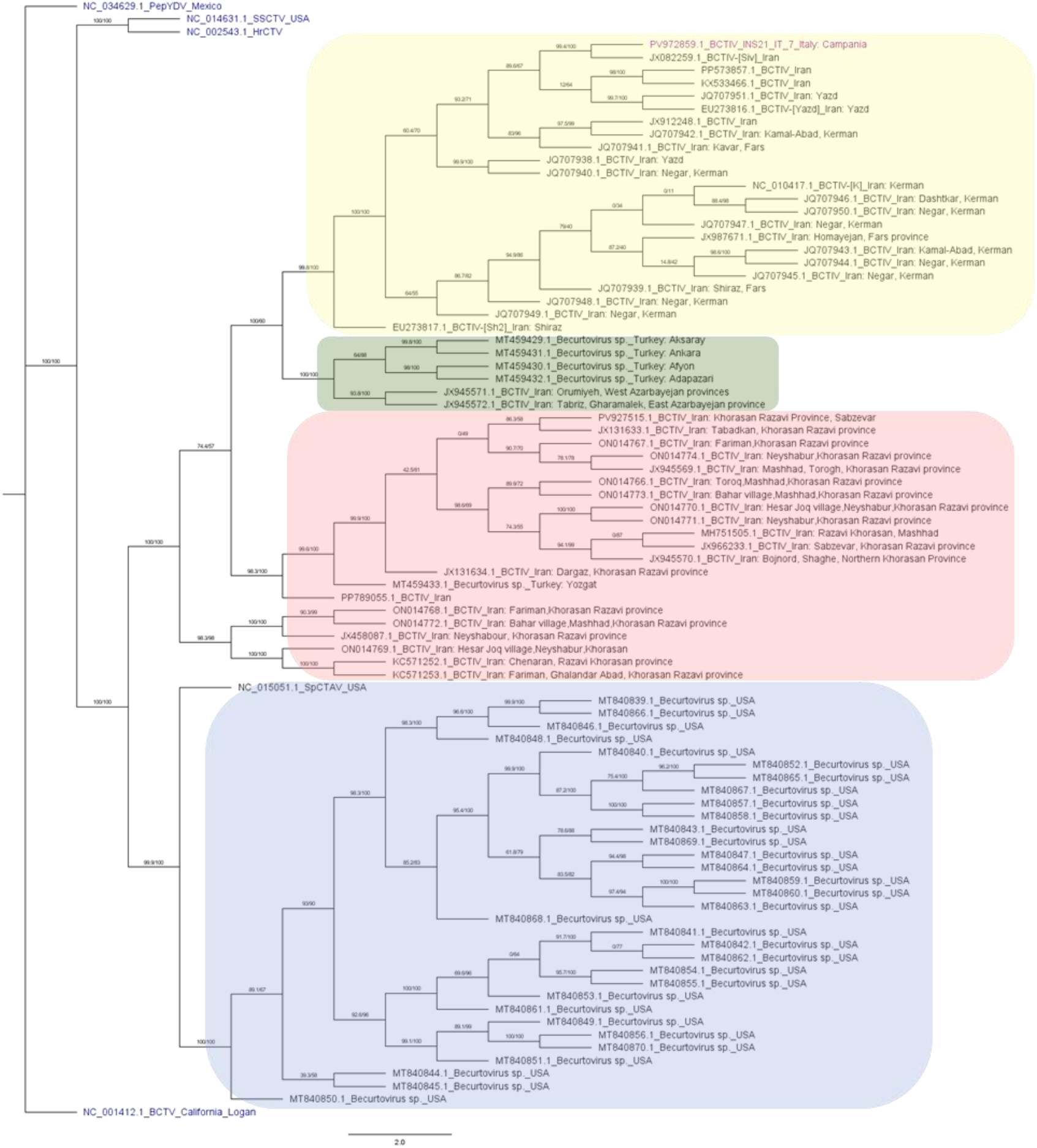
Maximum likelihood phylogenetic tree inferred using IQ-TREE based on the complete genome sequences of viral species belonging to the *Becurtovirus* genus, using *Curtovirus* reference genomes as outliers. Node values represent ultrafast bootstrap supports (UFBoot) calculated from 1,000 replicates, with values ≥70% considered well supported. The best-fit substitution model (TIM2+F+G4) was automatically selected by ModelFinder. The BCTIV complete genome sequenced in this work is shown in purple. NC_015051.1 SpCTAV indicates the *Becurtovirus* spinach curly top Arizona virus. The outliers, represented by the *Curtovirus* references genomes (NC_0014612.1 BCTV, beet curly top virus; NC_002543.1 HrCTV, horseradish curly top virus; NC_014631.1 SSCTV, spinach severe curly top virus; NC_034629.1 PepYDV, pepper yellow dwarf virus) are shown in blue. *Becurtovirus* clades corresponding to different geographical areas are highlighted with different colors: yellow indicates sequences from South-central Iran, green denotes sequences from Turkey, pink from Northwest Iran, and blue from USA.

The presence of BCTIV in cicadellids was confirmed by PCR using the individual cicadellid samples present in the three positive pools. Overall, 19 out of 31 samples (61%), all collected in June 2021, were PCR-positive (Table 1); in detail, BCTIV was detected in 4 out of 10, 8 out of 10, and 7 out of 11 samples present in the pools INS21_IT_6, 7 and 8, respectively (Fig. 2a). Two representative amplified fragments were Sanger-sequenced, confirming BCTIV-specific amplification.

**Figure 2.**
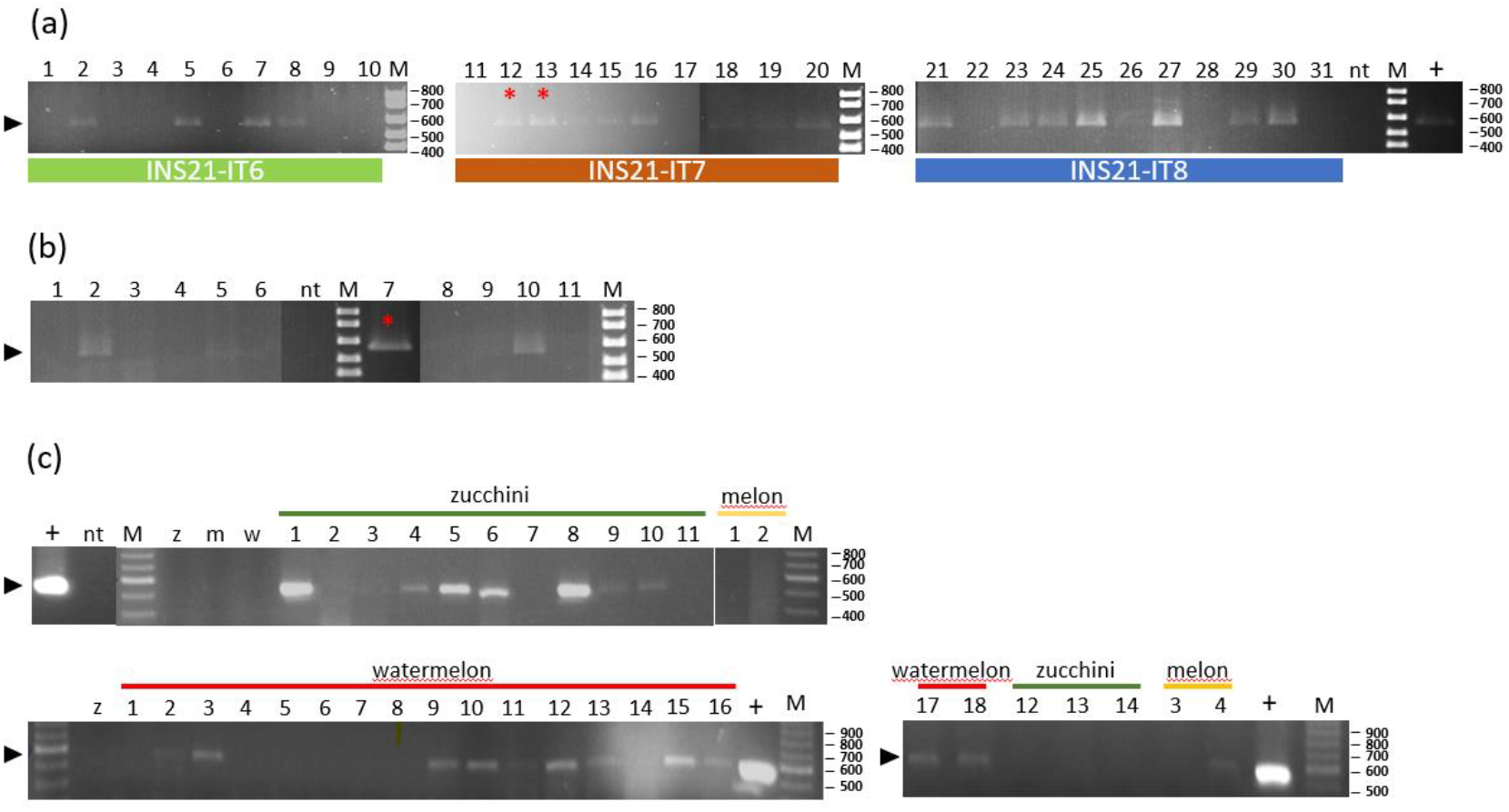
PCR validations using BCTIV-specific primers. (a) Analysis of 31 cicadellid samples forming the insect pools INS21_IT6, 7 and 8. (b) Analysis of 11 plant samples collected in/nearby fields where BCTIV-positive cicadellids were found. Lanes 1, 150621_w_1; 2, 150621_z_1; 3, 150621_z_2; 4, 150621_z_4; 5, 150621_z_5; 6, 150621_z_6; 7, 150621_w_3; 8, 160621_z_1; 9, 160621_z_2; 10, 160621_w_1; 11, 160621_w_2, listed in Table 1. (c) Analysis of seedlings artificially inoculated with the BCTIV agroclone: 14 zucchini, 18 watermelon, and 4 melon seedlings, tested 40 days post inoculation. The arrow indicates the BCTIV 559bp amplified fragment; M, 100bp marker; +, BCTIV positive control; nt, no template; z/m/w, zucchini, melon and watermelon negative control samples. Red asterisk indicates Sanger-sequenced amplicons.

The BCTIV-positive leafhopper samples were collected mainly in the Salerno area and, to a lesser extent, in Naples and Caserta areas. Although the survey was conducted across three years in different growing seasons, only cicadellids collected in early summer (June 2021) were BCTIV-positive, suggesting that the collection period might be relevant, possibly linked to the life cycle of the insects (Table 1; Fig. 3).

**Figure 3.**
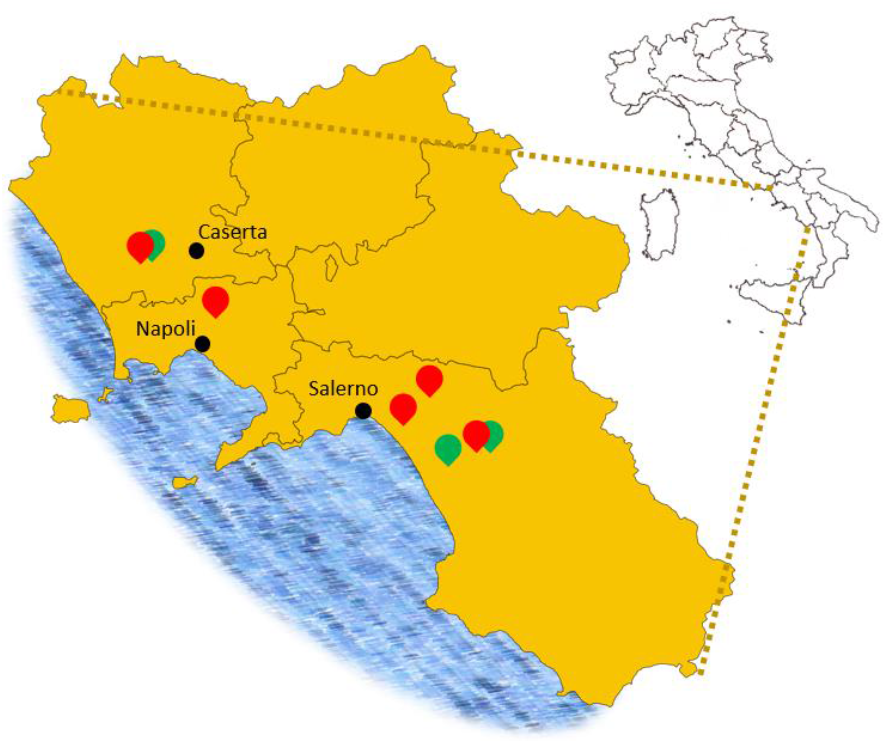
Geographical distribution of BCTIV-positive samples in the Campania region, Italy. Red and green markers indicate the sites where BCTIV-positive insect samples and plant pools were identified, respectively. Image modified from Wikimedia Commons. Original authors: Vonvikken and Marcomob, Public Domain.

Following the detection of BCTIV in insects collected in some cucurbit fields, we investigated the plants growing in and close to the same fields, where symptoms presumably linked to BCTIV infection were observed (Fig. 4 a-d). The presence of BCTIV was confirmed by PCR in 2 zucchini and 2 watermelon fields out of eleven tested (Table 1; Fig. 2b) and BCTIV-specific amplification was confirmed on selected amplified fragments by Sanger sequencing. Despite the relatively low infection rate found in the fields tested, our results highlight the urgent need for further searches in this region to assess the real distribution of this virus, concentrating on the major known BCTIV hosts, *i*.*e. Beta vulgaris* and *Spinacia oleracea*.

**Fig 4.**
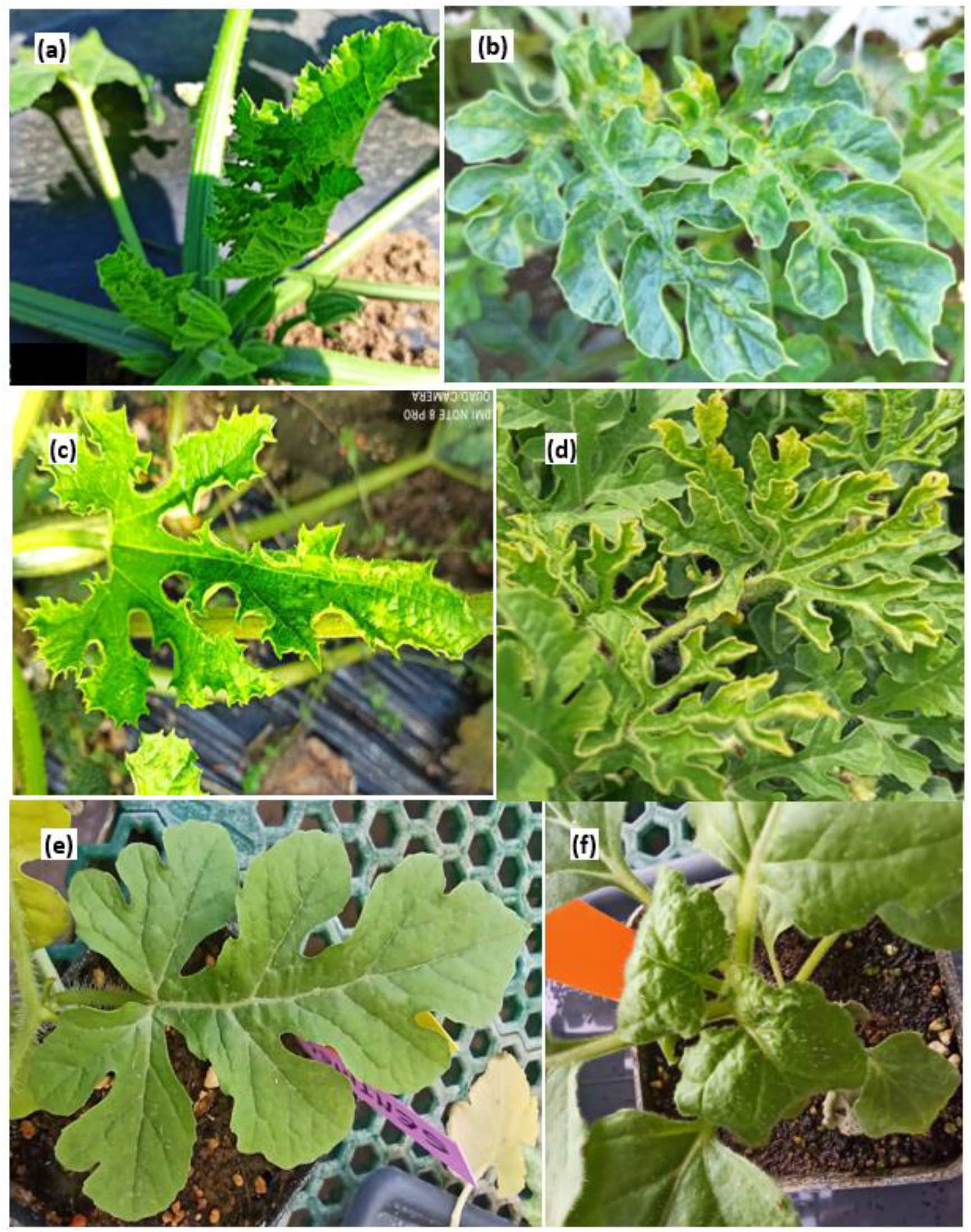
Symptoms attributable to BCTIV infection in field cultivated plants (panels A-D) and in experimentally inoculated plants (panels E and F). Yellow mosaic and curling symptoms on (A and C) young zucchini plants collected in fields 150621_z_1 and 150621_z_5 respectively; yellow mosaic and curling symptoms on (B and D) watermelon plants collected in field 150621_w_3 and 160621_w_1 (see also Table 1). (E) Symptoms on an adult leaf of a BCTIV-agroinoculated watermelon seedling (var. Sentinel, 40 dpi). (F) Typical crumpling symptoms on *N. benthamiana* young leaves at 7 dpi, used as positive control.

### Cicadellid analysis

Morphological examination of the cicadellids present in BCTIV-positive pools did not allow us to identify specimens of the genus *Neoaliturus* (*Circulifer*), which includes the species *N. (Circulifer*) *haematoceps*, the only known natural vector of BCTIV, previously recorded in the Mediterranean areas, including Italy [32]. Consistently, no Illumina-derived contigs from the tested cicadellids could be clearly assigned to this genus, based on the COI sequences available in the BOLD reference database. To further investigate the occurrence of *Neoaliturus* spp. individuals, a COI barcoding approach was adopted. Similarities ranging between 90.4-99.7% were detected with the genera *Exitianus* and *Psammotettix* belonging to the Cicadellidae family and similarity of 98,9% was found with the genus *Laodelphax* within the Delphacidae family. In two cases, sequence alignment was consistent only at the Cixiidae family level. Importantly, none of the amplified sequences reached at least 97% identity with the *Neoaliturus* COI gene accession present in the GenBank reference databases (Table 2).

**Table 2.**
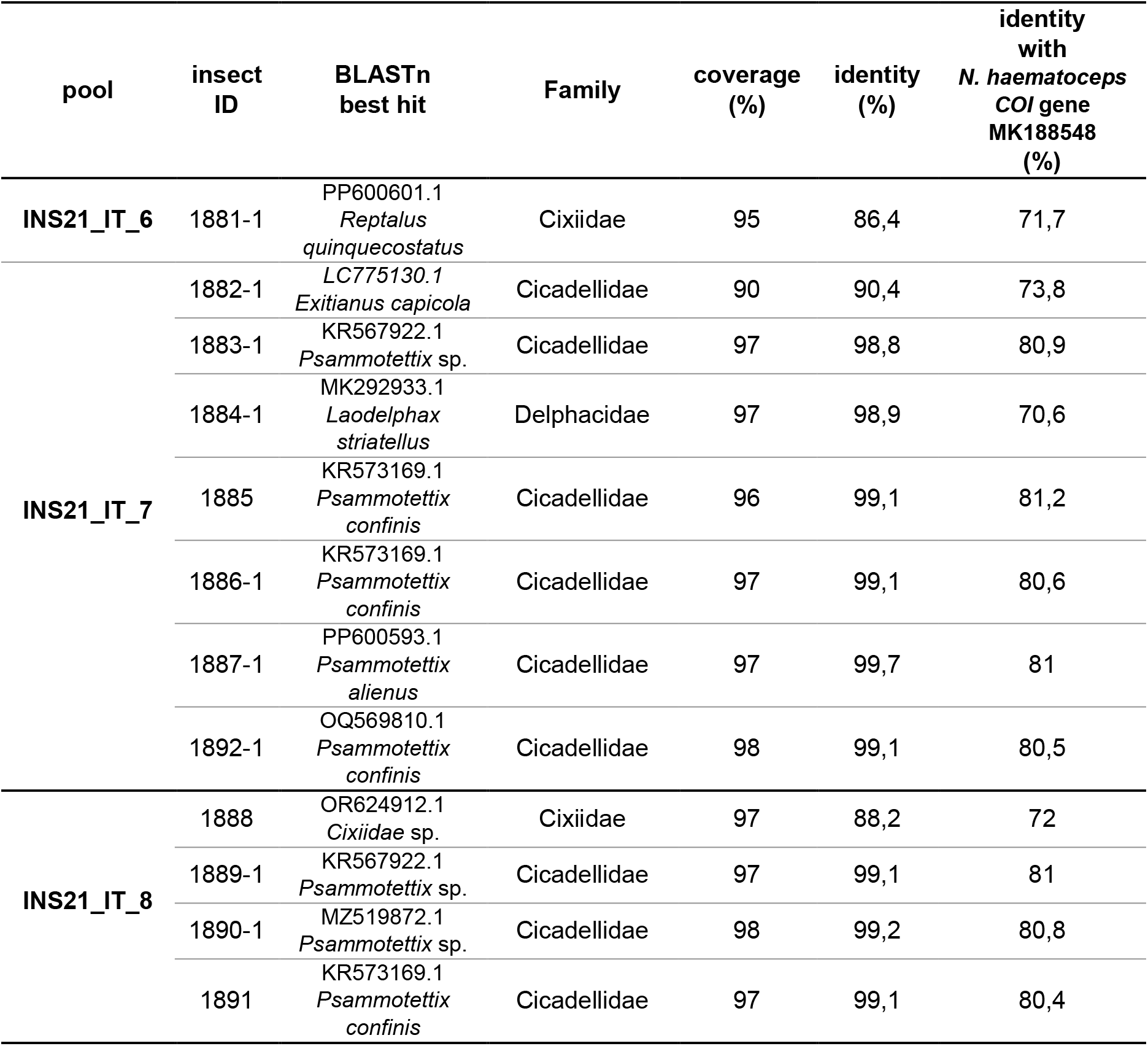
Results of barcoding analysis of individual insects present in BCTIV-positive pools. The best hit obtained by BLASTn analysis of the amplified COI sequence are shown, together with the coverage and identity percentages. The levels of identity with the reference COI gene of the *Neoaliturus haematoceps* are also indicated, the coverage percentage of the twelve COI sequences with the reference ranges from 92 to 97%. We considered the identity > 97% sufficient for genus assignment [33, 34]

Since the presence of *Neoaliturus* spp. could not be demonstrated in our samples by morphological or sequence analyses [33, 34], it is possible that this vector is rare in the environment and that the tested cicadellids only occasionally ingested BCTIV virions during feeding. Alternatively, we cannot exclude the presence of a previously unrecognized BCTIV vector active in the Mediterranean environment among the collected positive cicadellids, potentially representing an additional risk factor for virus establishment and spread.

Moreover, the lack of an updated integrative taxonomic framework for *Neoaliturus* emphasizes the need for further studies.

### Expanding the BCTIV host range

As cucurbits have not yet been clearly established as natural hosts of BCTIV [5, 6], to substantiate our findings, artificial inoculation of watermelon, melon and zucchini seedlings with the BCTIV infectious clone was performed. At 40 dpi, 11 out of 18 watermelon, 1 out of 4 melon, and 7 out of 11 zucchini seedlings tested positive in PCR (Fig. 2c). At this time point, watermelons showed reduced growth, chlorosis, greyish veins and light curling of leaves, accompanied by small necrotic lesions adjacent to veins (Fig. 4e), while chlorotic symptoms occurred on zucchini and melon plants. All *N. benthamiana* plants used as control showed typical BCTIV symptoms already at 7 dpi (Fig. 4f).

Altogether, the agroinoculation experiments and the molecular analysis of field samples support the conclusion that watermelon and zucchini represent new hosts for BCTIV and confirm that melon can host BCTIV, as recently reported [35]. Indeed, although previous studies reported that zucchini plants collected in Iran tested positive in ELISA using a curly top virus-specific antibody, no clear identification of the infecting virus was available at that time [36].

## 8. CONCLUDING REMARKS

The emergence of plant viruses in new areas involves ecological adaptation processes resulting from changes in virus™host and virus-vector associations over time and space. During a surveillance project in the Mediterranean area focused on tomato and cucurbit crops, we adopted an RCA-VEM approach on associated insects to investigate the circulating geminivirome. This strategy led us to identify for the first time in Europe and specifically in Italy the presence of BCTIV, a becurtovirus to date described only in the Asian continent (Iran and Anatolia) [7, 37, 38].

Overall, this study confirms that RCA-VEM is a powerful approach to conduct epidemiological surveillance of circulating virome, providing prompt alerts over a polyphagous and potentially invasive geminivirus. This diagnostic strategy might be adopted to monitor the introduction into new areas of other dangerous geminiviruses [39, 40] and associated satellites [41, 42], as well as their vectors. Such surveillance is especially crucial for Southern European countries, naturally exposed to phytopathological threats from warmer tropical and subtropical countries.

## 9. Author statements

### 9.1. Author contributions

L.M: data analysis, sample collection, review and editing. S.R: data analysis, laboratory analyses, review and editing. F.F: laboratory analyses. M.M: laboratory analyses. F.N: conceptualization, sample collection, laboratory analyses. UB: conceptualization, sample collection. D.M: sample collection, laboratory analyses. S.B: laboratory analyses, review and editing. M.B: sampling organization. G.P.A: sample collection, review and editing. A.M.V: funding acquisition, conceptualization, writing the original draft, laboratory analyses, review and editing. E.N: conceptualization, review and editing.

### 9.4. Ethical approval

Not applicable

### 9.5. Consent for publication

Not applicable

## 9.6. Acknowledgements

We wish to thank Elena Zocca and Luca Bordone for greenhouse work management; Simona Gargiulo, Flavia de Benedetta, and Roberta Ascolese for field samplings and Fortuna Miele for the field and laboratory activities.

