## Supplementary for "Vector-enabled metagenomics reveals the first detection of the geminivirus beet curly top Iran virus in Europe"

### Supplementary Figure 1

Maximum likelihood phylogenetic tree inferred using IQ-TREE based on the coat protein (CP) aminoacid sequences of viral species belonging to the Becurtovirus genus, using CP Curtovirus as outliers. Node values represent ultrafast bootstrap supports (UFBoot) calculated from 1,000 replicates, with values ≥70% considered well supported. The best-fit substitution model (JTT+G4) was automatically selected by ModelFinder. The BCTIV CP from genome sequenced in this work is shown in purple. YP_0042079231.1 SpCTAV indicates the Becurtovirus spinach curly top Arizona virus. The outliers, represented by the Curtovirus references genomes (NP_040559.1 BCTV, beet curly top virus; NP_066183.1 HrCTV, horseradish curly top virus; YP_003966135.1 SSCTV, spinach severe curly top virus; YP_009362975.1 PepYDV, pepper yellow dwarf virus) are shown in blue. Becurtovirus clades corresponding to different geographical areas are highlighted with different colors: the yellow rectangle indicates sequences from South-central Iran, the green one denotes sequences from Turkey, pink from Northwest Iran, and blue from USA.

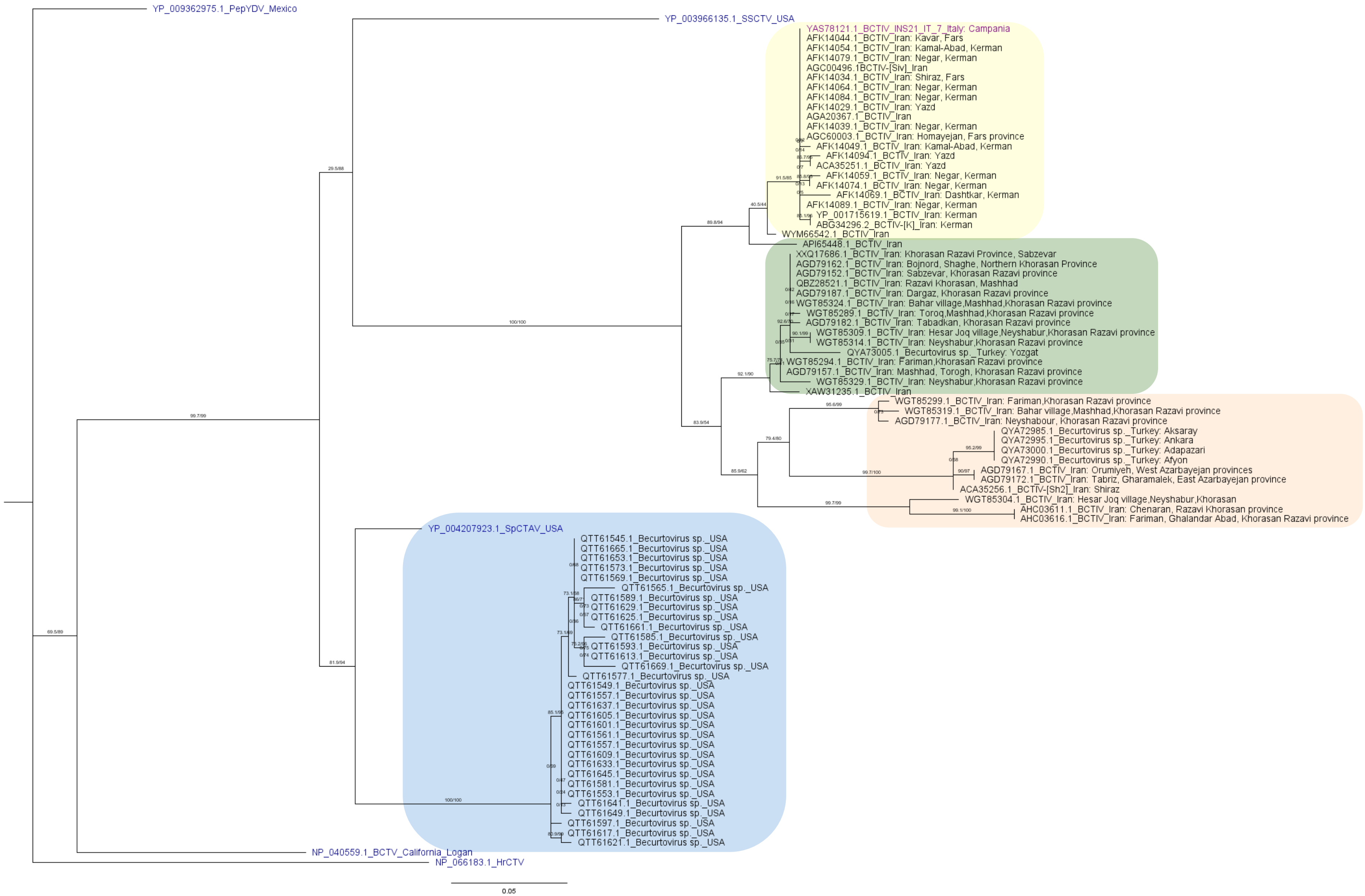

Supplementary Table 1 - Summary statistics of bioinformatics analysis of the insect pools

| Sample | Raw reads | Clean reads | Metaspades scaffolds | CAP3 contigs | Cap3 > 500nt | Reads mapping on BCTIV (NC_010417.1) | Coverage | Mean depth |
|---|---|---|---|---|---|---|---|---|
| INS20_IT1 | 128.154.968 | 115.904.632 | 769.924 | 67.917 | 67.519 | 0 | 0 | 0 |
| INS21_IT6 | 148.726.680 | 139.985.934 | 545.723 | 456.125 | 101.196 | 24 | 24.5 | 0.98x |
| INS21_IT7 | 160.173.312 | 150.180.006 | 1.343.474 | 1.226.065 | 385.100 | 242 | 100 | 11.94x |
| INS21_IT8 | 127.001.948 | 99.328.568 | 1.101.896 | 1.005.808 | 368.216 | 408 | 100 | 20.73x |
| INS21_IT13 | 132.138.936 | 118.653.166 | 1.056.837 | 958.023 | 304.588 | 0 | 0 | 0 |
| INS22_IT15 | 141.031.056 | 120.432.032 | 1.189.013 | 1.024.646 | 294.230 | 0 | 0 | 0 |
